## Supplementary material for "The enpp4 ectonucleotidase regulates kidney patterning signalling networks in *Xenopus* embryos": Supplmental data

This PDF file includes: Supplementary Fig. 1 to 7 and Supplementary Tables 1 to 5

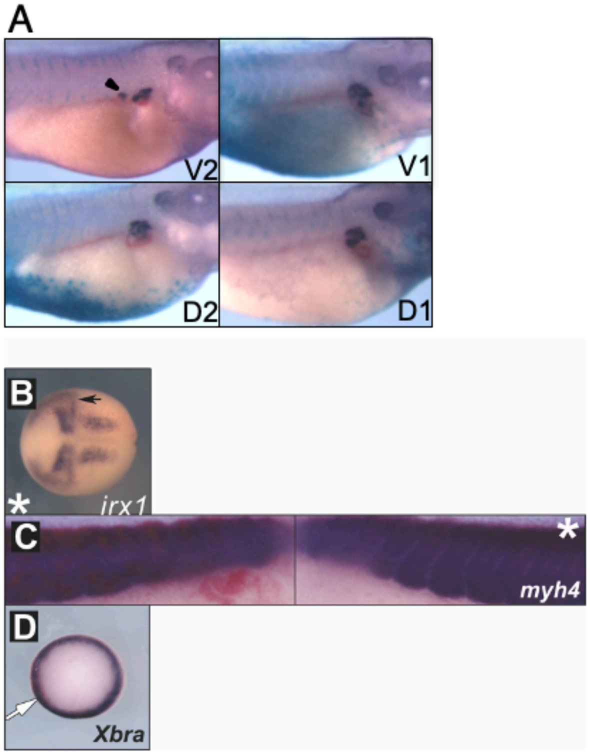


**Supplementary Fig. 1. Phenotypes caused by enpp4 mRNA injection (related to Fig. 1).** (A) Ectopic pronephric tissues were only induced following *enpp4* mRNA injection into the V2 blastomere. Embryos injected with 2ng of *enpp4* and 250 pg of *LacZ* mRNAs into the V2, V1, D2 or D1 blastomere at 8-cell stage were examined by whole mount antibody staining 3G8/4A6 at stage 41. The black arrowhead indicates the ectopic 3G8 staining. (B- D) No change in *irx1* expression and in mesoderm formation caused by enpp4 overexpression. Embryos targeted with 2ng of *enpp4* and 250 pg of *LacZ* mRNAs were examined by whole mount *in situ* hybridization with the *irx1* (B), *myf4* (C) and *xbra* (D) probes at stages 14, 32 and 10.5 respectively. The asterisk denotes the uninjected side of each embryo. The black arrow in B indicates the pronephros. The white arrow in D indicates the injection site marked by the red-gal staining. The raw data and statistical analyses are provided in Supplementary Table 1.

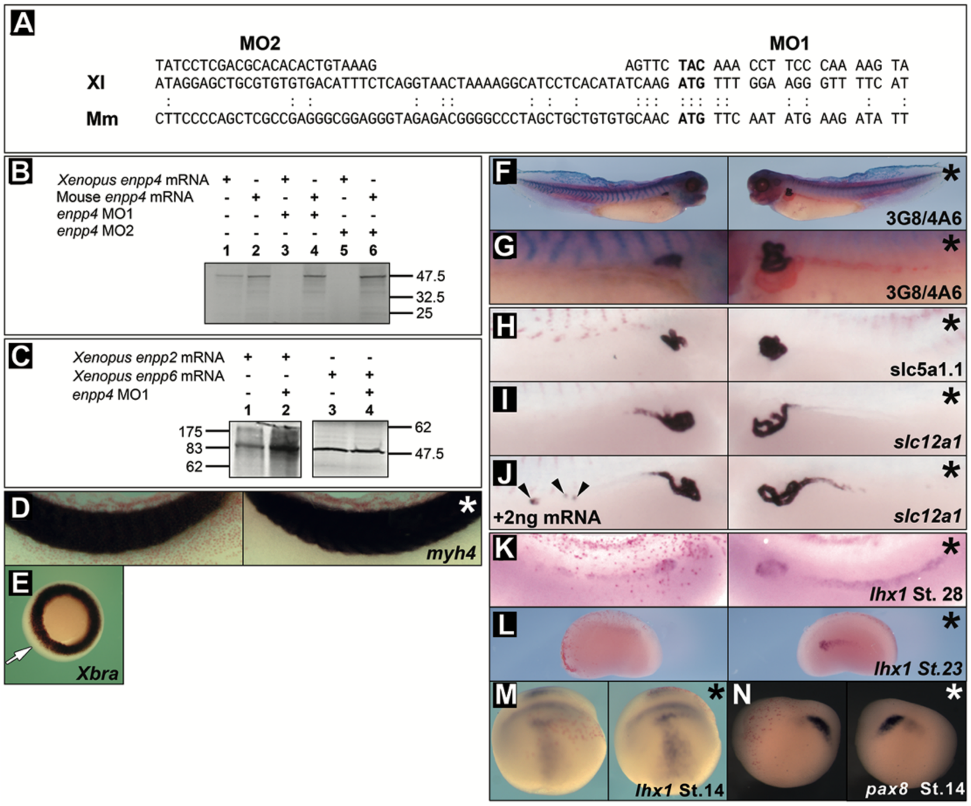


**Supplementary Fig. 2. *Enpp4* MO1 and MO2 specificity and phenotypes caused by injection of *enpp4* MO2 (related to Fig. 2).** (A-C) *enpp4* MO1 and 2 specifically inhibited *Xenopus* enpp4 translation. (A) Alignment of the 5'UTR of *Xenopus* (Xl) and mouse (Mm) *enpp4* sequences and position of *enpp4* MO1 and 2 in relation to *Xenopus enpp4* cDNA The ATG is indicated in bold and identical nucleotides (nt) by dots. Only 12 nt of *enpp4* MO1 and 3 nt of *enpp4* MO2 (located in the 5' UTR sequence) are conserved between the two species. (B) *Xenopus enpp4* but not mouse *enpp4* translation was blocked by *enpp4* MO1 and 2. Autoradiograph of a 10% SDS-PAGE gel of *in vitro* translated $^{35}$S-Methionine radiolabeled enpp4 proteins. Capped synthetic *enpp4* RNA was translated *in vitro* in the Rabbit Reticulocyte Lysate System (Promega) according to manufacturer's protocol. Translation of *Xenopus enpp4* mRNA (0.5 µg) (Lane 1) and mouse Enpp4 (Lane 2) produced a protein of 51 kDa. Translation of *Xenopus enpp4* mRNA was almost abolished in presence of 10 µg of *enpp4* MO1 or MO2 (Lane 3 and 5) while translation of mouse *Enpp4* was unaffected by *enpp4* MO1 or MO2 (Lane 4 and 6). (C) *enpp4* MO1 is specific to *enpp4* mRNA. Autoradiograph of 10% SDS-PAGE gel of *in vitro* translated $^{35}$S-Methionine radiolabeled *Xenopus* enpp2 and 6 proteins. *Xenopus enpp2 and 6* RNAs were translated *in vitro* in the Rabbit Reticulocyte Lysate System (Promega). Translation of *Xenopus enpp2* mRNA (0.5 µg) produced a protein of 99 kDa (Lane 1) that is unaffected by the addition of 10 µg *enpp4* MO1 (Lane 2). Translation of *Xenopus enpp6* mRNA (0.5 µg) produced a protein of 51 kDa (Lane 3) that is unaffected by the *enpp4* MO1 (Lane 4). Mm: *Mus musculus*; Xl: *Xenopus laevis.* (D-E) Microinjection of *enpp4* MO1 did not alter general mesoderm formation. Embryos injected with 10 ng of *enpp4* MO1 at 8-cell stage were examined by whole mount *in situ* hybridization with the *myf4* (D) and *xbra* (E) probes at stage 32 and at stage 10.5 respectively. Asterisk denotes uninjected sides. The arrow in (E) indicates the site of injection marked by red-gal staining. (F-N) Microinjection of *enpp4* MO2 resulted in similar phenotypes than those with *enpp4* MO1. Embryos injected with 10 ng of *enpp4* MO2 at 8-cell stage were examined by 3G8/4A6 antibody staining at stage 40 (F and G) or whole mount *in situ* hybridization with the following probes: *slc5a1.1* (H), *slc12a1* (I and J) at stage 37/38; *lhx1* at stage 28, 23 and 14 (K, L and M respectively) and *pax8* at stage 14 (N). Embryo shown in (J) was co-injected with 2 ng of mouse *Enpp4* mRNA for rescue experiment. Asterisk denotes uninjected sides. Arrowheads indicate ectopic *slc12a1* staining. The raw data and statistical analyses are provided in Supplementary Table 2.

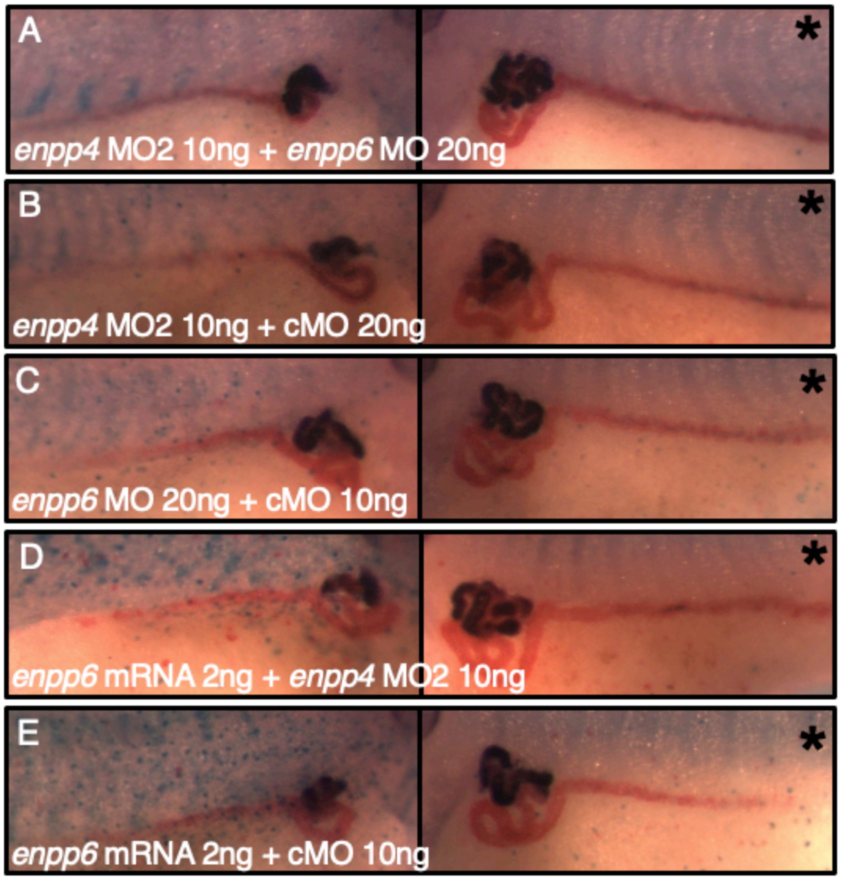


**Supplementary Fig. 3. Phenotypes caused by the co-injection of the *enpp4* and *enpp6* MOs or *enpp4* MO and *enpp6* mRNA (related to Fig. 2).** Injection of *enpp6* mRNA, but not *enpp6* MO, worsened the pronephric phenotypes caused by enpp4 knock-down. Injected embryos were analysed by 3G8/4A6 antibody staining at stage 41. (A) Embryo injected with 10 ng of *enpp4* MO2 and 20 ng of *enpp6* MO. (B) Embryo injected with 10 ng of *enpp4* MO2 and 20 ng of control MO. (C) Embryo injected with 20 ng of *enpp6* MO and 10 ng of control MO. (D) Embryo injected with 2 ng of *enpp6* mRNA and 10 ng of *enpp4* MO. (E) Embryo injected with 2 ng of *enpp6* RNA and 10 ng of control MO. Asterisk denotes uninjected sides. The raw data and statistical analyses are provided in Supplementary Table 2.

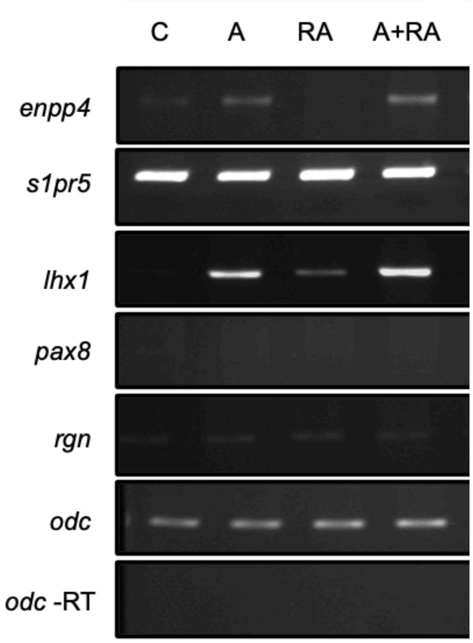


**Supplementary Fig. 4. Enpp4 expressionis regulated by activin but not by retinoic acid in animal cap assay (related to Fig. 3).** Animal cap were taken at blastula stage and cultured during 3 hours in the presence of activin (A), retinoic acid (RA) or activin + retinoic acid (A+RA) and total RNA extracted. RT-PCR was performed on treated and control (C) animal caps. The amplification of *odc* on samples with no reverse transcriptase (-RT) was carried out to control the purity of the RNAs.

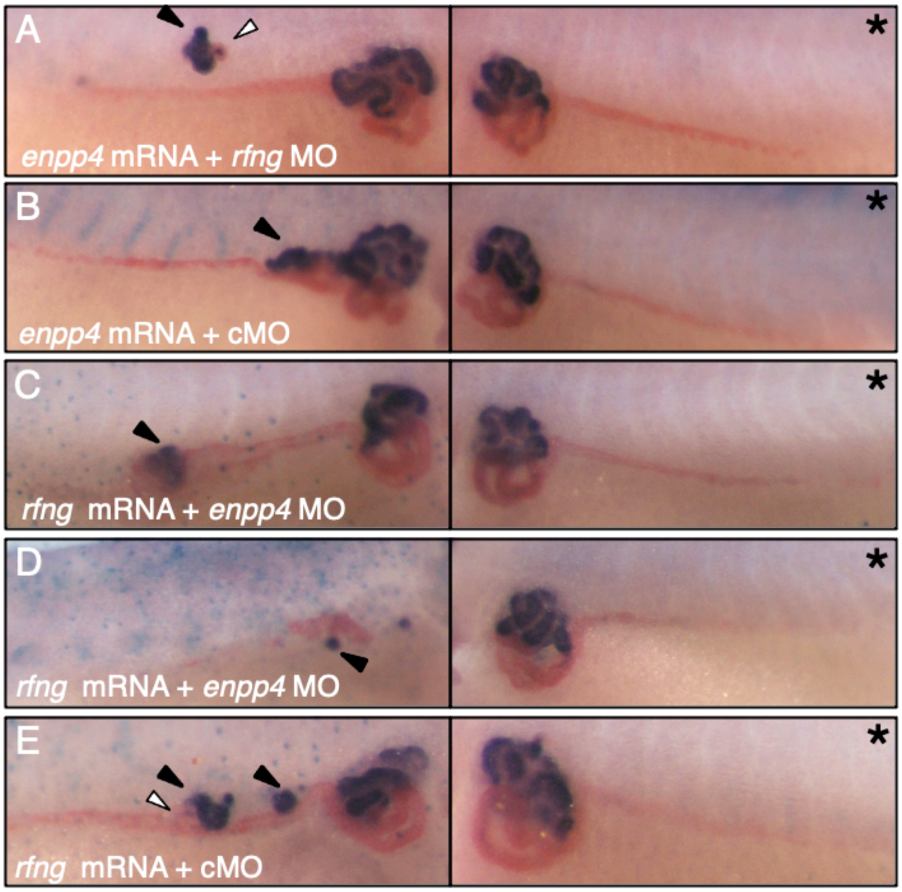


**Supplementary Fig. 5. Phenotypes caused by co-injection of *enpp4 or rfng* mRNA and MO (related to Fig. 4).** Injection of *rfng* MO did not prevent ectopic pronephros formation caused by enpp4 over-expression whereas co-injection of *enpp4* MO and *rfng* mRNA altered ectopic and endogenous pronephros formation. Injected embryos were harvested at stage 40, and analyzed by 3G8/4A6 antibody staining. (A) Embryo injected with 2 ng of *enpp4* mRNA and 20 ng of *rfng* MO. (B) Embryo injected with 2 ng of *enpp4* mRNA and 20 ng of control MO. (C, D) Embryo injected with 2 ng of *rfng* mRNA and 10 ng of *enpp*4 MO2. (E) Embryo injected 2 ng of *rfng* mRNA and 10 ng of control MO. Asterisk denotes uninjected sides. Black arrowheads indicate 3G8 ectopic staining and white arrowheads 4A6 ectopic staining. The raw data and statistical analyses are provided in Supplementary Table 3.

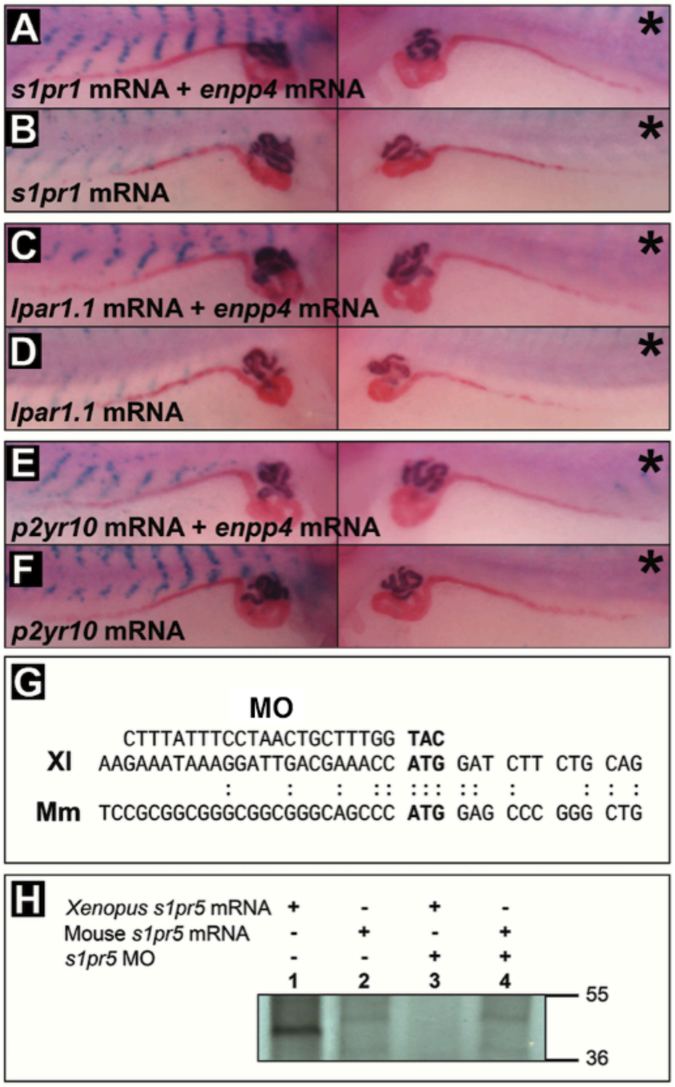


**Supplementary Fig. 6. Phenotypes caused by injection of *s1pr1, lpar1.1 and* p2yr10 mRNA and *in vitro* test of the s*1pr5.L* MO specificity (related to Fig. 6).** (A-F) Microinjection of *s1pr1, lpar1.1* and *p2yr10* mRNA did not induce ectopic pronephros even when co-injected with *enpp*4 mRNA. Injected embryos were harvested at stage 40, and analyzed by 3G8/4A6 antibody staining. (A) Embryo injected with 2 ng of *s1pr1* mRNA and 1ng of *enpp*4 mRNA or (B) 2ng of *s1pr1* mRNA alone. (C) Embryo injected with 2ng of *lpar1.1* mRNA and 1ng of *enpp*4 mRNA or (D) 2ng of *lpar1.1* mRNA alone. (E) Embryo injected with 2ng of *p2yr10* mRNA and 1ng of *enpp*4 mRNA or (F) with 2ng of *p2yr10* mRNA alone. Asterisk denotes uninjected sides. The raw data and statistical analyses are provided in Supplementary Table 5. (G-H) *s1pr5.L* MO specifically inhibited *Xenopus s1pr5.L* mRNA translation. (G) Alignment of the 5'UTR of *Xenopus* (Xl) and mouse (Mm) *s1pr5* sequences and position of *s1pr5.L* MO in relation to *Xenopus s1pr5.L* cDNA. The ATG is indicated in bold and identical nucleotides (nt) by dots. This alignment shows that only 8 nt of *s1pr5.L* MO sequence are conserved between the two species. (H) *Xenopus s1pr5.L* but not mouse *S1pr5* translation was blocked by *s1pr5.L* MO; autoradiograph of a 10% SDS-PAGE gel of *in vitro* translated $^{35}$S-Methionine radiolabeled s1pr5 proteins. Capped synthetic *s1pr5* RNAs was translated *in vitro* in the Rabbit Reticulocyte Lysate System (Promega) according to manufacturer's protocol. Lane 1, Translation of *Xenopus s1pr5.L* mRNA (1µg) produced a protein of 43 kDa. Lane 2, Translation of mouse *S1pr5* mRNA (1 µg) produced a protein of 42 kDa. Lane 3, Translation of *Xenopus s1pr5.L* mRNA was severely affected by the addition of 10 µg of *s1pr5.L* MO. Lane 4, Translation of mouse *S1pr5* was unaffected by *s1pr5*.L MO. Mm: *Mus musculus*; Xl: *Xenopus laevis.*

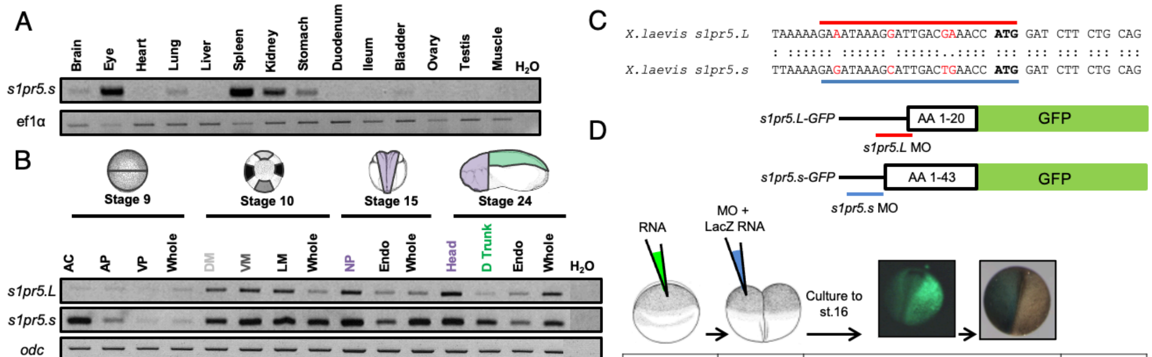


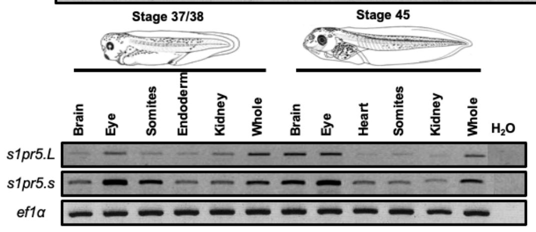


| RNA injected | MO injected | Scored embryos | | | MO efficacity (%) |
|---|---|---|---|---|---|
| | | GFP⁺ | One-side GFP⁺ | Opposite side LacZ⁺ | |
| *s1pr5.L-GFP* | *s1pr5.L* MO | 17 | 15 | 11 | 64.7 |
| | *s1pr5.s* MO | 15 | 8 | 5 | 33.3 |
| | cMO | 16 | 4 | 2 | 12.5 |
| *s1pr5.s-GFP* | *s1pr5.L* MO | 22 | 14 | 9 | 40.9 |
| | *s1pr5.s* MO | 15 | 10 | 10 | 66.7 |
| | cMO | 19 | 10 | 2 | 10.5 |
| *GFP* | *s1pr5.L* MO | 21 | 7 | 1 | 5.8 |
| | *s1pr5.s* MO | 17 | 4 | 2 | 11.8 |
| | cMO | 16 | 4 | 2 | 12.5 |

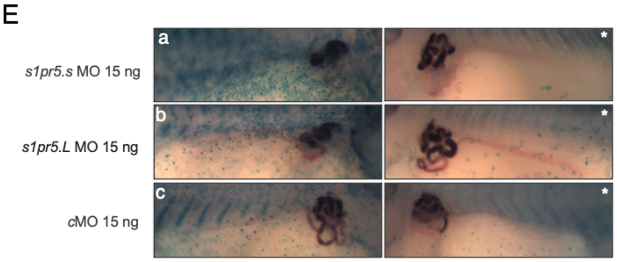


**Supplementary Fig. 7. Expression profile of the *sp1r5.S* receptor gene, *in vivo* test of MO specificity and phenotypes caused by its knock-down (related to Fig. 6).** (A) The *s1pr5.S* receptor displays a restricted expression profile in the adult frog and is expressed in the mesonephric tissue. Adult tissues were dissected and total RNAs from adult tissues were extracted using Trizol (Invitrogen) following the manufacturer's protocol. RT-PCR was performed along with negative control. (B) The novel *s1pr5.S* receptor is expressed in the kidney with an expression profile similar to its homelog, the *s1pr5.L* receptor. *X.laevis* embryos were dissected and total RNA extracted. RT-PCR was performed on dissected tissues and control whole embryos along with negative control. *S1pr5.S* was amplified using the following primers (forward primer, 5'- ggaggtctcgttgctgtctc -3' and the reverse primer, 5'- cggtagctcctcgtgtaacc -3') with the annealing temperature of 53°C and 32 cycles, Each RT-PCR product was sequenced to confirm the specificity of the amplifications. (C) Alignment of the 5'UTR of *X.laevis s1pr*5 homeologs sequences and position of *s1pr5.L and s1pr5.S* MOs. The ATG is indicated in red and identical nucleotides by dots. Only 4 nucleotides over the whole MOs sequences differ between the two genes. (D) Efficacy of the *s1pr5.L* and *s1pr5.L* MOs. Schematic representation of the GFP fusion proteins containing the 5'UTR and part of the coding region of *s1pr5.L* and *s1pr5.S*. The position of the *s1pr5.L* and *s1pr5.s* MO are indicated. *Xenopus* embryos were injected with the *s1pr5.LGFP*, *s1pr5.S -GFP* and *GFP* mRNAs at one cell stage followed by unilateral injections of *s1pr5.L, s1pr5.S* and control MOs in presence of *LacZ* mRNA at 2-cell stage. GFP⁺ positive embryos were sorted at stage 16 and stained for ß-galactosidase activity. The numbers of embryos GFP+ fluorescent on one side and LacZ⁺ on the opposite side and the percentage of efficiency of the different MOs to inhibit *in vivo* the translation of the different mRNAs are indicated in the table. (E) Microinjection of *s1pr5.S* MO induces a similar kidney phenotype than the microinjection of the *s1pr5.L* MO. Injected embryos analysed by 3G8/4A6 antibody staining at stage 41. (a) Embryo injected with 15ng of *s1pr5.S* MO (b) Embryo injected with 15ng of *s1pr5.L* MO. (c) Embryo injected with 15ng of control MO. Asterisk denotes uninjected sides. The raw data and statistical analyses are provided in Supplementary Table 5.

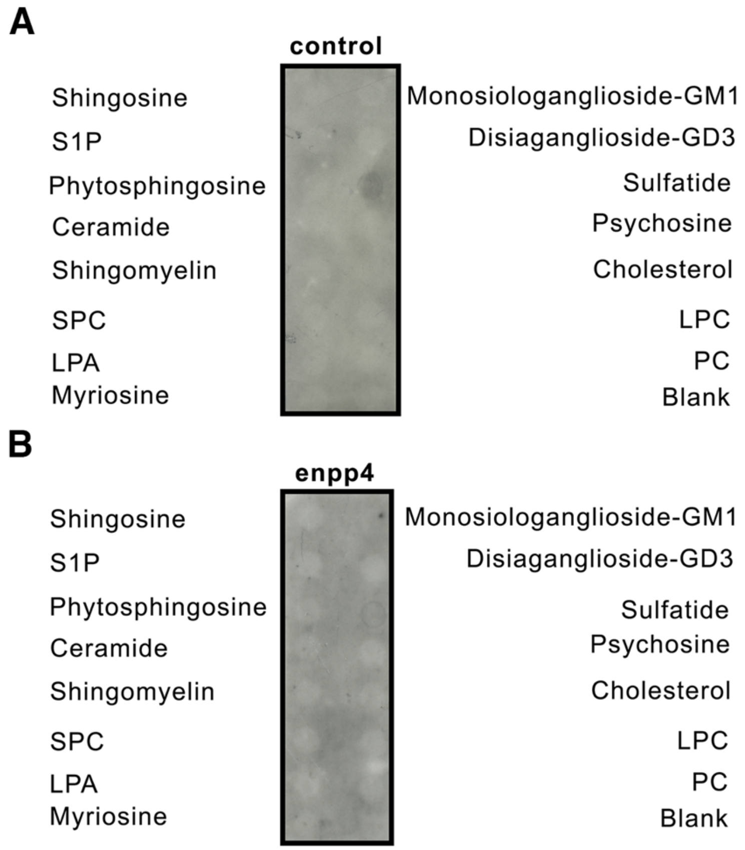


**Supplementary Fig. 8. Enpp4 does not interact with bioactive lipids metabolized by enpp2, enpp6 and enpp7 (related to Fig. 7).** ShingoStrips$^{TM}$ was incubated with membrane protein extracts from (A) control pcDNA3.1 transfected CHO cells or (B) from enpp4 over-expressing CHO cells and the bound enpp4 protein detected with anti-Xlenpp4 serum. LPA, lysophosphatidic acid; LPC, lysophosphocholine; PC, phospatidylcholine; S1P, shingosine-1-phosphate; SPC, Shingosylphosphorylcholine.

### Supplementary Table 1A

| Histological Analysis | Figure | Injection | Marker analysed | Normal | Enlarged | Reduced | Absent | Ectopic | Total number of scored embryos |
|---|---|---|---|---|---|---|---|---|---|
| | | | | | | Phenotypes | | | |
| 1 | 1B-E | *enpp4* mRNA 2ng | 3G8 | 47 (52%) | 16 (18%) | 7 (8%) | 0 (0%) | 21 (23%) | 91 |
| 2 | 1B-E | *enpp4* mRNA 2ng | 4A6 | 63 (69%) | 18 (20%) | 8 (9%) | 0 (0%) | 2 (2%) | 91 |
| 3 | S1A | *enpp4 mRNA 2ng V2* | 3G8 | 19 (47.5%) | 8 (20%) | 5 (12.5%) | 0 (0%) | 8 (20%) | 40 |
| 4 | S1A | *enpp4 mRNA 2ng V2* | 4A6 | 29 (72.5%) | 6 (15%) | 4 (10%) | 0 (0%) | 1 (2.5%) | 40 |
| 5 | S1A | *enpp4 mRNA 2ng V1* | 3G8 | 30 (88%) | 0 (0%) | 4 (12%) | 0 (0%) | 0 (0%) | 34 |
| 6 | S1A | *enpp4 mRNA 2ng V1* | 4A6 | 30 (88%) | 0 (0%) | 4 (12%) | 0 (0%) | 0 (0%) | 34 |
| 7 | S1A | *enpp4 mRNA 2ng D2* | 3G8 | 25 (96%) | 0 (0%) | 1 (4%) | 0 (0%) | 0 (0%) | 26 |
| 8 | S1A | *enpp4 mRNA 2ng D2* | 4A6 | 24 (92%) | 0 (0%) | 2 (8%) | 0 (0%) | 0 (0%) | 26 |
| 9 | S1A | *enpp4 mRNA 2ng D1* | 3G8 | 22 (92%) | 1 (4%) | 1 (4%) | 0 (0%) | 0 (0%) | 24 |
| 10 | S1A | *enpp4 mRNA 2ng D1* | 4A6 | 23 (96%) | 0 (0%) | 1 (4%) | 0 (0%) | 0 (0%) | 24 |
| 11 | 1M | Mouse *enpp4* mRNA 2ng | 3G8 | 19 (30%) | 0 (0%) | 8 (13%) | 1 (2%) | 35 (56%) | 63 |
| 12 | 1M | Mouse *enpp4* mRNA 2ng | 4A6 | 32 (51%) | 12 (19%) | 11 (17.5%) | 1 (2%) | 7 (11%) | 63 |
| 13 | 1N | *enpp4* T72S mRNA 2ng | 3G8 | 50 (76%) | 1 (1.5%) | 15 (23%) | 0 (0%) | 0 (0%) | 66 |
| 14 | 1N | *enpp4* T72S mRNA 2ng | 4A6 | 52 (79%) | 0 (0%) | 14 (21%) | 0 (0%) | 0 (0%) | 66 |
| 15 | | *enpp4* T72A mRNA 2ng | 3G8 | 42 (68%) | 1 (2%) | 18 (29%) | 0 (0%) | 1 (1%) | 62 |
| 16 | | *enpp4* T72A mRNA 2ng | 4A6 | 51 (82%) | 0 (0%) | 11 (18%) | 0 (0%) | 0 (0%) | 62 |
| 17 | 1O | *enpp4* D36N mRNA 2ng | 3G8 | 53 (77%) | 6 (9%) | 9 (13%) | 0 (0%) | 1 (1%) | 69 |
| 18 | 1O | *enpp4* D36N mRNA 2ng | 4A6 | 62 (90%) | 1 (1%) | 6 (9%) | 0 (0%) | 0 (0%) | 69 |
| 19 | | *enpp4* D189N mRNA 2ng | 3G8 | 57 (75%) | 2 (3%) | 8 (10.5%) | 4 (5%) | 5 (7%) | 76 |
| 20 | | *enpp4* D189N mRNA 2ng | 4A6 | 68 (89.5%) | 1 (1%) | 4 (5%) | 3 (4%) | 0 (0%) | 76 |
| 21 | | *LacZ* mRNA 250pg | 3G8 | 52 (91%) | 0 (0%) | 5 (9%) | 0 (0%) | 0 (0%) | 57 |
| 22 | | *LacZ* mRNA 250pg | 4A6 | 53 (93%) | 0 (0%) | 4 (7%) | 0 (0%) | 0 (0%) | 57 |
| 23 | 1P | *enpp4* mRNA 2ng | *slc5a1.1* | 27 (47%) | 8 (14%) | 5 (9%) | 0 (0%) | 17 (30%) | 57 |
| 24 | | *LacZ* mRNA 250pg | *slc5a1.1* | 60 (94%) | 0 (0%) | 3 (5%) | 1 (2%) | 0 (0%) | 64 |
| 25 | 1Q | *enpp4* mRNA 2ng | *slc12a1* | 30 (57%) | 14 (25%) | 3 (5%) | 0 (0%) | 10(17.5%) | 57 |
| 26 | | *LacZ* mRNA 250pg | *slc12a1* | 63 (98%) | 0 (0%) | 1 (2%) | 0 (0%) | 0 (0%) | 64 |
| 27 | 1R | *enpp4* mRNA 2ng | *clcnkb* | 40 (69%) | 11 (19%) | 7 (12%) | 0 (0%) | 0 (0%) | 58 |
| 28 | | *LacZ* mRNA 250pg | *clcnkb* | 60 (94%) | 0 (0%) | 2 (3%) | 2 (3%) | 0 (0%) | 64 |
| 29 | 1S | *enpp4* mRNA 2ng | *gata3* | 17 (39.5%) | 1 (2%) | 25 (58%) | 0 (0%) | 0 (0%) | 43 |
| 30 | | *LacZ* mRNA 250pg | *gata3* | 33 (89%) | 0 (0%) | 4 (11%) | 0 (0%) | 0 (0%) | 37 |
| 31 | 1T | *enpp4* mRNA 2ng | *wt1* | 26 (44%) | 20 (34%) | 13 (22%) | 0 (0%) | 0 (0%) | 59 |
| 32 | | *LacZ* mRNA 250pg | *wt1* | 84 (95.5%) | 1 (1%) | 3 (3%) | 0 (0%) | 0 (0%) | 88 |
| 33 | 1U | *enpp4* mRNA 2ng | *nphs1* | 32 (36%) | 22 (25%) | 30 (34%) | 1 (1%) | 3 (3%) | 88 |
| 34 | | *LacZ* mRNA 250pg | *nphs1* | 52 (96%) | 0 (0%) | 2 (4%) | 0 (0%) | 0 (0%) | 54 |
| 35 | 1V | *enpp4* mRNA 2ng | *lhx1* | 20 (39%) | 31 (61%) | 0 (0%) | 0 (0%) | 0 (0%) | 51 |
| 36 | | *LacZ* mRNA 250pg | *lhx1* | 76 (96%) | 1 (1%) | 2 (2%) | 0 (0%) | 0 (0%) | 79 |
| 37 | 1W | *enpp4* mRNA 2ng | *pax8* | 14 (28%) | 35 (70%) | 0 (0%) | 0 (0%) | 1 (2%) | 50 |
| 38 | | *LacZ* mRNA 250pg | *pax8* | 72 (97%) | 0 (0%) | 2 (3%) | 0 (0%) | 0 (0%) | 74 |
| 39 | 1X | *enpp4 mRNA 2ng* | *lhx1* | 30 (65%) | 9 (20%) | 7 (15%) | 0 (0%) | 0 (0%) | 46 |
| 40 | | *LacZ mRNA 250pg* | *lhx1* | 28 (67%) | 4 (9%) | 10 (24%) | 0 (0%) | 0 (0%) | 42 |
| 41 | 1Y | *enpp4 mRNA 2ng* | *pax8* | 20 (29%) | 12 (17%) | 24 (34%) | 0 (0%) | 15 (21%) | 70 |
| 42 | | *LacZ mRNA 250pg* | *pax8* | 34 (60%) | 2 (3%) | 21 (37%) | 0 (0%) | 0 (0%) | 57 |
| 43 | S1C | *enpp4 mRNA 2ng* | *irx1* | 24(73%) | 2 (6%) | 7 (21%) | 0 (0%) | 0 (0%) | 33 |

| Histological Analysis | Figure | Injection | Marker analysed | Phenotypes<br>Normal | Enlarged | Reduced | Absent | Ectopic | Total number of scored embryos |
|---|---|---|---|---|---|---|---|---|---|
| 44 | | *LacZ RNA 250 pg* | *irx1* | 20 (95%) | 0 (0%) | 1 (5%) | 0 (0%) | 0 (0%) | 21 |
| 45 | S1D | *enpp4* mRNA 2ng | *myh4* | 55 (100%) | 0 (0%) | 0 (0%) | 0 (0%) | 0 (0%) | 55 |
| 46 | | *LacZ* mRNA 250pg | *myh4* | 53 (98%) | 0 (0%) | 0 (0%) | 0 (0%) | 1 (2%) | 54 |
| 47 | S1E | *enpp4* mRNA 2ng | *xbra* | 39 (85%) | 0 (0%) | 7 (15%) | 0 (0%) | 0 (0%) | 46 |
| 48 | | *LacZ* mRNA 250pg | *xbra* | 46 (88.5%) | 0 (0%) | 6 (11.5%) | 0 (0%) | 0 (0%) | 52 |

**Supplementary Table 1B**

| Histological analysis compared | Figures compared | Injections compared | Marker | Calculated p-value | Calculated p-value (with Bonferroni correction) | Standardized p-value (with Bonferroni correction) |
|---|---|---|---|---|---|---|
| 1/21 | 1B | *enpp4/LacZ* | 3G8 | 9.53E-09 | 1.19E-06 | (0;0.001] |
| 2/22 | 1B | *enpp4/LacZ* | 4A6 | 1.57E-04 | 1.96E-02 | (0.01;0.05) |
| 11/21 | 1M | *Enpp4(mouse)/LacZ* | 3G8 | 1.53E-14 | 1.91E-12 | (0;0.001] |
| 12/22 | 1M | *Enpp4(mouse)/LacZ* | 4A6 | 3.15E-07 | 3.94E-05 | (0;0.001] |
| 5/1 | 1N/1C | *enpp4T72S/enpp4* | 3G8 | 6.04E-09 | 7.55E-07 | (0;0.001] |
| 6/2 | 1N/1C | *enpp4T72S/enpp4* | 4A6 | 2.10E-05 | 2.62E-03 | (0.001;0.01) |
| 7/1 | | *enpp4T72A/ enpp4* | 3G8 | 5.89E-08 | 7.36E-06 | (0;0.001] |
| 8/2 | | *enpp4T72A/ enpp4* | 4A6 | 7.79E-05 | 9.74E-03 | (0.001;0.01) |
| 9/1 | 1O/1C | *enpp4D36N/ enpp4* | 3G8 | 3.00E-05 | 3.76E-03 | (0.001;0.01) |
| 10/2 | 1O/1C | *enpp4D36N/ enpp4* | 4A6 | 4.83E-04 | 6.04E-02 | NS |
| 11/1 | | *enpp4D189N/ enpp4* | 3G8 | 2.01E-05 | 2.51E-03 | (0.001;0.01) |
| 12/2 | | *enpp4D189N/ enpp4* | 4A6 | 4.10E-05 | 5.13E-03 | (0.001;0.01) |
| 15/16 | 1P | *enpp4/LacZ* | *slc5a1.1* | 6.36E-11 | 7.95E-09 | (0;0.001] |
| 17/18 | 1Q | *enpp4/LacZ* | *sclc12a1* | 6.24E-11 | 7.81E-09 | (0;0.001] |
| 19/20 | 1R | *enpp4/LacZ* | *clcnkb* | 1.06E-05 | 1.33E-03 | (0.001;0.01) |
| 21/22 | 1S | *enpp4/LacZ* | *gata3* | 6.04E-06 | 7.55E-04 | (0;0.001] |
| 23/24 | 1T | *enpp4/LacZ* | *wt1* | 3.89E-13 | 4.87E-11 | (0;0.001] |
| 25/26 | 1U | *enpp4/LacZ* | *nphs1* | 9.66E-13 | 1.21E-10 | (0;0.001] |
| 27/28 | 1V | *enpp4/LacZ* | *lhx1* (st28) | 2.37E-15 | 2.96E-13 | (0;0.001] |
| 29/30 | 1W | *enpp4/LacZ* | *pax8* (st28) | 4.88E-20 | 6.10E-18 | (0;0.001] |
| 31/32 | 1X | *enpp4/LacZ* | *lhx1* (neurula) | 3.37E-01 | 1.00E+00 | NS |
| 33/34 | 1Y | *enpp4/LacZ* | *pax8* (neurula) | 3.13E-06 | 3.91E-04 | (0;0.001] |
| 43/41 | S1A | *enpp4V1/enpp4V2* | *3G8* | 6.57E-05 | 8.21E-03 | (0.001;0.01) |
| 44/42 | S1A | *enpp4V1/enpp4V2* | *4A6* | 5.79E-02 | 1.00E+00 | NS |
| 45/41 | S1A | *enpp4D2/enpp4V2* | *3G8* | 1.47E-04 | 1.84E-02 | (0.01;0.05) |
| 46/42 | S1A | *enpp4D2/enpp4V2* | *4A6* | 1.31E-01 | 1.00E+00 | NS |
| 47/41 | S1A | *enpp4D1/enpp4V2* | *3G8* | 2.31E-03 | 2.89E-01 | NS |
| 48/42 | S1A | *enpp4D1/enpp4V2* | *4A6* | 9.21E-02 | 1.00E+00 | NS |
| 35/36 | S1C | *enpp4/LacZ* | *irx1* | 1.00E-01 | 1.00E+00 | NS |
| 37/38 | S1D | *enpp4/LacZ* | *myh4* | 4.95E-01 | 1.00E+00 | NS |
| 39/40 | S1E | *enpp4/LacZ* | *xbra* | 7.67E-01 | 1.00E+00 | NS |

**Supplementary Table 1. Scoring analysis of kidney phenotypes in embryos over-expressing enpp4 (related to Fig.1 and Fig. S1). (S1A) Results from immunohistochemistry and *in situ* hybridization of injected embryos.** Embryos were injected with *enpp4* wild type or mutant RNA. Embryos were scored for differences between the injected side (identified by Blue or Red Gal staining) and uninjected side acting as contra-lateral control side. The numbers of embryos displaying renal phenotype are indicated. The corresponding percentages are indicated in bracket. Each histological analysis was numbered as reference for the statistical comparisons in Supplementary Table 1B. Only the pronephros phenotype on the injected side is indicated. **(S1B) Statistical analyses.** Statistical comparisons between pairs of histological analysis listed in Supplementary Table 1A were carried out as indicated in the table. The Bonferroni multiple testing correction was applied to all Fisher's Exact Test. The calculated p value before and after correction and a standardized corrected p-value are given. NS: not significant.

### Supplementary Table 2A

| Histological analysis | Figure | Injection | Marker analysed | Phenotypes: Normal | Enlarged | Reduced | Absent | Ectopic | Total number of scored embryos |
|---|---|---|---|---|---|---|---|---|---|
| 1 | 2A. B | *enpp4* MO1 10ng | 3G8 | 37 (35%) | 0 (0%) | 70 (65%) | 0 (0%) | 0 (0%) | 107 |
| 2 | 2A. B | *enpp4* MO1 10ng | 4A6 | 67 (63%) | 0 (0%) | 30 (28%) | 10 (9%) | 0 (0%) | 107 |
| 3 | | cMO 10ng | 3G8 | 27 (100%) | 0 (0%) | 0 (0%) | 0 (0%) | 0 (0%) | 27 |
| 4 | | cMO 10ng | 4A6 | 27 (100%) | 0 (0%) | 0 (0%) | 0 (0%) | 0 (0%) | 27 |
| 5 | 2C | *enpp4 MO1 10ng+MO2 10ng* | 3G8 | 2 (10%) | 0 (0%) | 18 (90%) | 0 (0%) | 0 (0%) | 20 |
| 6 | 2C | *enpp4 MO1 10ng+MO2 10ng* | 4A6 | 0 (0%) | 0 (0%) | 15 (75%) | 5 (25%) | 0 (0%) | 20 |
| 7 | | *enpp4 MO1 10ng + cMO 10ng* | 3G8 | 5 (33%) | 0 (0%) | 10 (67%) | 0 (0%) | 0 (0%) | 15 |
| 8 | | *enpp4 MO1 10ng + cMO 10ng* | 4A6 | 7 (47%) | 0 (0%) | 8 (53%) | 0 (0%) | 0 (0%) | 15 |
| 9 | | *enpp4 MO2 10ng + cMO 10ng* | 3G8 | 9 (56%) | 0 (0%) | 7 (44%) | 0 (0%) | 0 (0%) | 16 |
| 10 | | *enpp4 MO2 10ng + cMO 10ng* | 4A6 | 7 (44%) | 0 (0%) | 9 (56%) | 0 (0%) | 0 (0%) | 16 |
| 11 | | *cMO 20ng* | 3G8 | 10 (59%) | 0 (0%) | 7 (41%) | 0 (0%) | 0 (0%) | 17 |
| 12 | | *cMO 20ng* | 4A6 | 10 (53%) | 0 (0%) | 7 (47%) | 0 (0%) | 0 (0%) | 17 |
| 13 | 2D | *enpp4* MO1 10ng | *slc5a1.1* | 27 (42%) | 0 (0%) | 37 (58%) | 0 (0%) | 0 (0%) | 64 |
| 14 | | cMO 10ng | *slc5a1.1* | 34 (100%) | 0 (0%) | 0 (0%) | 0 (0%) | 0 (0%) | 34 |
| 15 | 2E | *enpp4* MO1 10ng | *slc12a1* | 33 (44%) | 0 (0%) | 42 (56%) | 0 (0%) | 0 (0%) | 75 |
| 16 | 2F | *enpp4* MO1 10ng + *enpp4* mRNA 2ng | *slc12a1* | 47 (65%) | 0 (0%) | 17 (24%) | 0 (0%) | 8 (11%) | 72 |
| 17 | | cMO 10ng | *slc12a1* | 39 (83%) | 0 (0%) | 8 (17%) | 0 (0%) | 0 (0%) | 47 |
| 18 | 2G | *enpp4* MO1 10ng | *clcnkb* | 19(47.5%) | 0 (0%) | 21 (53.5%) | 0 (0%) | 0 (0%) | 40 |
| 19 | | cMO 10ng | *clcnkb* | 42 (76%) | 0 (0%) | 13 (24%) | 0 (0%) | 0 (0%) | 55 |
| 20 | 2H | *enpp4* MO1 10ng | *gata3* | 29 (69%) | 0 (0%) | 13 (31%) | 0 (0%) | 0 (0%) | 42 |
| 21 | | cMO 10ng | *gata3* | 33 (89%) | 0 (0%) | 4 (11%) | 0 (0%) | 0 (0%) | 37 |
| 22 | 2I | *enpp4* MO1 10ng | *wt1* | 32 (94%) | 1 (3%) | 1 (3%) | 0 (0%) | 0 (0%) | 34 |
| 23 | | cMO 10ng | *wt1* | 32 (94%) | 0 (0%) | 2 (6%) | 0 (0%) | 0 (0%) | 34 |
| 24 | 2J | *enpp4* MO1 10ng | *nphs1* | 34 (83%) | 1 (2%) | 6 (15%) | 0 (0%) | 0 (0%) | 41 |
| 25 | | cMO 10ng | *nphs1* | 45 (92%) | 1 (2%) | 3 (6%) | 0 (0%) | 0 (0%) | 49 |
| 26 | 2K | *enpp4* MO1 10ng | *lhx1* | 15 (65%) | 0 (0%) | 8 (35%) | 0 (0%) | 0 (0%) | 23 |
| 27 | | cMO 10ng | *lhx1* | 21 (100%) | 0 (0%) | 0 (0%) | 0 (0%) | 0 (0%) | 21 |
| 28 | 2L | *enpp4* MO1 10ng | *lhx1* | 5 (25%) | 0 (0%) | 15 (75%) | 0 (0%) | 0 (0%) | 20 |
| 29 | | cMO 10ng | *lhx1* | 14 (70%) | 0 (0%) | 6 (30%) | 0 (0%) | 0 (0%) | 20 |
| 30 | 2M | *enpp4 MO1 10ng + cMO 10ng* | *lhx1* | 14 (31%) | 0 (0%) | 17 (37%) | 15 (31%) | 0 (0%) | 46 |
| 31 | 2N | *enpp4 MO1 10ng+MO2 10ng* | *lhx1* | 5 (16%) | 2 (6%) | 18 (58%) | 6 (19%) | 0 (0%) | 31 |
| 32 | | *cMO 20ng* | *lhx1* | 30 (77%) | 4 (10%) | 5 (13%) | 0 (0%) | 0 (0%) | 39 |
| 33 | 2O | *enpp4 MO1 10ng + cMO 10ng* | *pax8* | 20 (29%) | 1 (1%) | 49 (70%) | 0 (0%) | 0 (0%) | 70 |
| 34 | | *cMO 20ng* | *pax8* | 17 (63%) | 4 (15%) | 6 (22%) | 0 (0%) | 0 (0%) | 27 |
| 35 | S2D | *enpp4* MO1 10ng | *myh4* | 47 (100%) | 0 (0%) | 0 (0%) | 0 (0%) | 0 (0%) | 47 |
| 36 | | cMO 10ng | *myh4* | 59 (100%) | 0 (0%) | 0 (0%) | 0 (0%) | 0 (0%) | 59 |
| 37 | S2E | *enpp4* MO1 10ng | *xbra* | 16 (84%) | 0 (0%) | 3 (16%) | 0 (0%) | 0 (0%) | 19 |
| 38 | | cMO 10ng | *xbra* | 20 (95%) | 0 (0%) | 1 (5%) | 0 (0%) | 0 (0%) | 21 |
| 39 | S2F.G | *enpp4* MO2 10ng | 3G8 | 44 (51%) | 0 (0%) | 43 (49%) | 0 (0%) | 0 (0%) | 87 |
| 40 | S2F.G | *enpp4* MO2 10ng | 4A6 | 66 (76%) | 0 (0%) | 21 (24%) | 0 (0%) | 0 (0%) | 87 |
| 41 | S2H | *enpp4* MO2 10ng | *slc5a1.1* | 26 (68%) | 0 (0%) | 12 (32%) | 0 (0%) | 0 (0%) | 38 |

| Histological analysis | Figure | Injection | Marker analysed | Phenotypes | | | | | Total number of scored embryos |
|---|---|---|---|---|---|---|---|---|---|
| | | | | Normal | Enlarged | Reduced | Absent | Ectopic | |
| 42 | S2I | *enpp4* MO2 10ng | *slc5a12a1* | 24 (86%) | 0 (0%) | 4 (14%) | 0 (0%) | 0 (0%) | 28 |
| 43 | S2J | *enpp4* MO2 10ng + *enpp4* mRNA 2ng | *slc5a12a1* | 14 (50%) | 0 (0%) | 0 (0%) | 0 (0%) | 14 (50%) | 28 |
| 44 | S2K | *enpp4* MO2 10ng | *lhx1* | 11 (55%) | 3 (15%) | 6 (30%) | 0 (0%) | 0 (0%) | 20 |
| 45 | S2L | *enpp4* MO2 10ng | *lhx1* | 9 (45%) | 0 (0%) | 11 (55%) | 0 (0%) | 0 (0%) | 20 |
| 46 | S2M | *enpp4 MO2 10ng + cMO 10ng* | *lhx1* | 19 (31%) | 3 (5%) | 39 (64%) | 0 (0%) | 0 (0%) | 61 |
| 47 | S2N | *enpp4 MO2 10ng + cMO 10ng* | *pax8* | 4 (21%) | 0 (0%) | 11 (58%) | 4 (21%) | 0 (0%) | 19 |
| 48 | S3A | *enpp4 MO2 10ng + enpp6 MO 20ng* | 3G8 | 25 (33%) | 0 (0%) | 49 (64%) | 2 (3%) | 0 (0%) | 76 |
| 49 | | *enpp4 MO2 10ng + enpp6 MO 20ng* | 4A6 | 28 (37%) | 0 (0%) | 47 (62%) | 1 (1%) | 0 (0%) | 76 |
| 50 | S3B | *enpp4 MO2 10ng +cMO 20ng* | 3G8 | 27 (40%) | 0 (0%) | 39 (62%) | 2 (3%) | 0 (0%) | 68 |
| 51 | | *enpp4 MO2 10ng +cMO 20ng* | 4A6 | 22 (32%) | 0 (0%) | 42 (62%) | 4 (6%) | 0 (0%) | 68 |
| 52 | S3C | *enpp6 MO 20ng +cMO 10ng* | 3G8 | 31 (47%) | 0 (0%) | 34 (51%) | 1 (2%) | 0 (0%) | 66 |
| 53 | | *enpp6 MO2 20ng +cMO 10ng* | 4A6 | 23 (35%) | 0 (0%) | 42 (63%) | 1 (2%) | 0 (0%) | 66 |
| 54 | | *cMO 30ng* | 3G8 | 44 (70%) | 0 (0%) | 19 (30%) | 0 (0%) | 0 (0%) | 63 |
| 55 | | *cMO 30ng* | 4A6 | 54 (86%) | 0 (0%) | 9 (14%) | 0 (0%) | 0 (0%) | 63 |
| 56 | S3D | *enpp6 mRNA 2ng + enpp4 MO2 10ng* | 3G8 | 14 (20%) | 0 (0%) | 45 (63%) | 12 (17%) | 0 (0%) | 71 |
| 57 | | *enpp6 mRNA 2ng + enpp4 MO2 10ng* | 4A6 | 24 (34%) | 0 (0%) | 28 (39%) | 19 (27%) | 0 (0%) | 71 |
| 58 | | *enpp4 MO2 10ng* | 3G8 | 23 (47%) | 0 (0%) | 25 (51%) | 1 (2%) | 0 (0%) | 49 |
| 59 | | *enpp4 MO2 10ng* | 4A6 | 28 (57%) | 0 (0%) | 21 (43%) | 0 (0%) | 0 (0%) | 49 |
| 60 | S3E | *enpp6 mRNA 2ng + cMO 10ng* | 3G8 | 1 (3%) | 0 (0%) | 27 (79%) | 6 (18%) | 0 (0%) | 34 |
| 61 | | *enpp6 mRNA 2ng + cMO 10ng* | 4A6 | 4 (12%) | 0 (0%) | 22 (65%) | 8 (23%) | 0 (0%) | 34 |

### Supplementary Table 2B

| Histological analysis compared | Figures compared | Injections compared | Marker | Calculated p-value | Calculated p-value (with Bonferroni correction) | Standardized p-value (with Bonferroni correction) |
|---|---|---|---|---|---|---|
| 1/3 | 2B | *enpp4 MO1/cMO* | 3G8 | 5.67E-11 | 7.09E-09 | (0;0.001] |
| 2/4 | 2B | *enpp4 MO1/ cMO* | 4A6 | 1.54E-04 | 1.92E-02 | (0.01;0.05] |
| 39/3 | S2G | *enpp4 MO2/ cMO* | 3G8 | 3.98E-07 | 4.97E-05 | (0;0.001] |
| 40/4 | S2G | *enpp4 MO2/ cMO* | 4A6 | 3.21E-03 | 4.01E-01 | NS |
| 5/7 | 2C/2B | *enpp4 MO1+2/MO1* | 3G8 | 1.12E-01 | 1.00E+00 | NS |
| 6/8 | 2C/2B | *enpp4 MO1+2/MO1* | 4A6 | 4.39E-04 | 5.48E-02 | NS |
| 5/9 | 2C/S2G | *enpp4 MO1+2/MO2* | 3G8 | 4.17E-03 | 5.21E-01 | NS |
| 6/10 | 2C/S2G | *enpp4 MO1+2/MO2* | 4A6 | 3.51E-04 | 4.39E-02 | (0.01;0.05] |
| 48/50 | S3A/S3B | *enpp6MO+enpp4 MO2/enpp4MO2* | 3G8 | 7.29E-01 | 1.00E+00 | NS |
| 49/51 | S3A/S3B | *enpp6MO+enpp4 MO2/enpp4MO2* | 4A6 | 3.21E-01 | 1.00E+00 | NS |
| 48/52 | S3A/3C | *enpp6MO+enpp4 MO2/enpp6MO* | 3G8 | 2.46E-01 | 1.00E+00 | NS |
| 49/53 | S3A/3C | *enpp6MO+enpp4 MO2/enpp6MO* | 4A6 | 9.30E-01 | 1.00E+00 | NS |
| 52/50 | S3C | *enpp6MO/enpp4MO2* | 3G8 | 6.10E-01 | 1.00E+00 | NS |
| 53/51 | S3C | *enpp6MO/enpp4MO2* | 4A6 | 5.13E-01 | 1.00E+00 | NS |
| 56/58 | S3D/S3B | *enpp6RNA+enpp4MO2/enpp4MO2* | 3G8 | 8.35E-04 | 1.04E-01 | NS |
| 57/59 | S3D/S3B | *enpp6RNA+enpp4MO2/enpp4MO2* | 4A6 | 3.32E-05 | 4.16E-03 | (0.001;0.01] |
| 56/60 | S3D/S3E | *enpp6RNA+enpp4MO2/enpp6RNA+cMO* | 3G8 | 5.58E-02 | 1.00E+00 | NS |
| 57/61 | S3D/S3E | *enpp6RNA+enpp4MO2/enpp6RNA+cMO* | 4A6 | 2.37E-02 | 1.00E+00 | (NS |
| 13/14 | 2D | *enpp4 MO1/cMO* | *slc5a1.1* | 1.26E-09 | 1.57E-07 | (0;0.001] |
| 41/14 | S2H | *enpp4 MO2/cMO* | *slc5a1.1* | 2.12E-04 | 2.65E-02 | (0.01;0.05] |
| 15/17 | 2E | *enpp4 MO1/cMO* | *sclc12a1* | 2.39E-05 | 2.98E-03 | (0.001;0.01] |
| 42/17 | S2I | *enpp4 MO2/cMO* | *sclc12a1* | 1.00E+00 | 1.00E+00 | NS |
| 16/15 | 2F/2E | *enpp4 MO1+RNA/ enpp4 MO1* | *sclc12a1* | 9.47E-06 | 1.18E-03 | (0.001;0.01] |
| 43/42 | S2J/S2H | *enpp4 MO2+RNA/ enpp4 MO2* | *sclc12a1* | 3.91E-06 | 4.89E-04 | (0;0.001] |
| 18/19 | 2G | *enpp4 MO1/cMO* | *clcnkb* | 4.96E-03 | 6.19E-01 | NS |
| 20/21 | 2H | *enpp4 MO1/cMO* | *gata3* | 5.28E-02 | 1.00E+00 | NS |
| 22/23 | 2I | *enpp4 MO1/cMO* | *wt1* | 1.00E+00 | 1.00E+00 | NS |
| 24/25 | 2J | *enpp4 MO1/cMO* | *nphs1* | 4.36E-01 | 1.00E+00 | NS |
| 26/27 | 2K | *enpp4 MO1/cMO* | *lhx1* (st28) | 3.91E-03 | 4.89E-01 | NS |
| *44/27* | S2K | *enpp4 MO2/cMO* | *lhx1* (st28) | 4.79E-04 | 5.99E-02 | NS |
| *28/29* | 2L | *enpp4 MO1/cMO* | *lhx1* (st22) | 1.04E-02 | 1.00E+00 | (NS |
| *45/29* | S2L | *enpp4 MO2/cMO* | *lhx1* (st22) | 2.00E-01 | 1.00E+00 | NS |
| *30/32* | 2M | *enpp4 MO1/cMO* | *lhx1* (neurula) | 5.95E-08 | 7.44E-06 | (0;0.001] |
| *46/32* | S2M | *enpp4 MO2/cMO* | *lhx1* (neurula) | 6.92E-07 | 8.65E-05 | (0;0.001] |
| *31/30* | 2N/2M | *enpp4 MO1+2/MO1* | *lhx1* (neurula) | 5.47E-02 | 1.00E+00 | NS |
| *31/46* | 2N/S2M | *enpp4 MO1+2/MO2* | *lhx1* (neurula) | 2.40E-03 | 3.00E-01 | NS |
| *33/34* | 2O | *enpp4 MO1/cMO* | *pax8*(neurula) | 2.23E-05 | 2.79E-03 | (0.001;0.01] |
| *47/34* | S2N | *enpp4 MO2/cMO* | *pax8*(neurula) | 3.31E-04 | 4.14E-02 | (0.01;0.05] |
| *35/36* | S2D | *enpp4 MO1/cMO* | myh4 | 1.00E+00 | 1.00E+00 | NS |
| *37/38* | S2E | *enpp4 MO1/cMO* | xbra | 3.31E-01 | 1.00E+00 | NS |

**Supplementary Table 2. Scoring analysis of kidney phenotypes in enpp4 morphants (related to Fig.2, Fig.S2 and S3). (S2A) Results from immunohistochemistry and in situ hybridization of injected embryos**. Embryos were injected with *enpp4* MO1 or MO2 alone or in combination. Embryos were scored for differences between the injected side (identified by Blue or Red Gal staining) and uninjected side acting as contra-lateral control side. The numbers of embryos displaying renal phenotype are indicated. The corresponding percentages are indicated in bracket. Each histological analysis was numbered as reference for the statistical comparisons in Supplementary Table 2B. Only the pronephros phenotype on the injected side is indicated. **(S2B) Statistical analyses.** Statistical comparisons between pairs of histological analysis listed in Supplementary Table 2A were carried out as indicated in the table. The Bonferroni multiple testing correction was applied to all Fisher's Exact Test. The calculated p value before and after correction and a standardized corrected p-value are given. NS: not significant.

### Supplementary Table 3A

| Histological Analysis | Figure | Injection | Marker stained | Phenotypes: Normal | Enlarged | Reduced | Absent | Ectopic | Total number of scored embryos |
|---|---|---|---|---|---|---|---|---|---|
| 1 | 3A | *enpp4* mRNA 2ng | *raldh1a2* | 12 (32%) | 8 (22%) | 4 (11%) | 0 (0%) | 13 (35%) | 37 |
| 2 | 3B | *enpp4* mRNA 2ng | *rdh10* | 18 (53%) | 12 (35%) | 0 (0%) | 0 (0%) | 4 (12%) | 34 |
| 3 | 3C | *enpp4* mRNA 2ng | *cyp26a1* | 43 (98%) | 1 (2%) | 0 (0%) | 0 (0%) | 0 (0%) | 44 |
| 4 | 3D | *enpp4* mRNA 2ng | *notch1* | 17 (23%) | 40 (53%) | 13 (17%) | 0 (0%) | 5 (7%) | 75 |
| 5 | 3E | *enpp4* mRNA 2ng | *dll1* | 19 (23%) | 19 (23%) | 7 (9%) | 3 (4%) | 32 (40%) | 81 |
| 6 | 3F | *enpp4* mRNA 2ng | *jag1* | 21 (26%) | 35 (44%) | 8 (10%) | 0 (0%) | 16 (20%) | 81 |
| 7 | 3G | *enpp4* mRNA 2ng | *wnt4* | 10 (24%) | 13 (32%) | 11 (27%) | 0 (0%) | 7 (17%) | 41 |
| 8 | 3H | *enpp4* MO1 10ng | *raldh1a2* | 25 (75%) | 0 (0%) | 8 (25%) | 0 (0%) | 0 (0%) | 33 |
| 9 | | cMO 10ng | *raldh1a2* | 34 (97%) | 0 (0%) | 1 (3%) | 0 (0%) | 0 (0%) | 35 |
| 10 | 3I | *enpp4* MO1 10ng | *rdh10* | 26 (79%) | 0 (0%) | 7 (21%) | 0 (0%) | 0 (0%) | 33 |
| 11 | | cMO 10ng | *rdh10* | 47(100%) | 0 (0%) | 0 (0%) | 0 (0%) | 0 (0%) | 47 |
| 12 | 3J | *enpp4* MO1 10ng | *cyp26a1* | 37 (88%) | 0 (0%) | 5 (12%) | 0 (0%) | 0 (0%) | 42 |
| 13 | | cMO 10ng | *cyp26a1* | 43 (93%) | 0 (0%) | 3 (7%) | 0 (0%) | 0 (0%) | 46 |
| 14 | 3K | *enpp4* MO1 10ng | *notch1* | 33 (80%) | 4 (10%) | 4 (10%) | 0 (0%) | 0 (0%) | 41 |
| 15 | | cMO 10ng | *notch1* | 39 (97%) | 1 (3%) | 0 (0%) | 0 (0%) | 0 (0%) | 40 |
| 16 | 3L | *enpp4* MO1 10ng | *dll1* | 31 (67%) | 0 (0%) | 15 (33%) | 0 (0%) | 0 (0%) | 46 |
| 17 | | cMO 10ng | *dll1* | 34 (85%) | 1 (3%) | 5 (12%) | 0 (0%) | 0 (0%) | 40 |
| 18 | 3M | *enpp4* MO1 10ng | *jag1* | 26 (62%) | 0 (0%) | 16 (38%) | 0 (0%) | 0 (0%) | 42 |
| 19 | | cMO 10ng | *jag1* | 39 (95%) | 0 (0%) | 2 (5%) | 0 (0%) | 0 (0%) | 41 |
| 20 | 3N | *enpp4* MO1 10ng | *wnt4* | 4 (10%) | 0 (0%) | 32 (82%) | 3 (8%) | 0 (0%) | 39 |
| 21 | | cMO 10ng | *wnt4* | 28 (85%) | 0 (0%) | 5 (15%) | 0 (0%) | 0 (0%) | 33 |
| 22 | S4A | *enpp4* mRNA 2ng + *rfng* MO 20ng | 3G8 | 9 (24%) | 1 (3%) | 12 (32%) | 0 (0%) | 15 (41%) | 37 |
| 23 | | *enpp4* mRNA 2ng +rfngMO 20ng | 4A6 | 20 (54%) | 0 (0%) | 12 (32%) | 0 (0%) | 5 (14%) | 37 |
| 24 | S4B | *enpp4* mRNA 2ng + cMO 20ng | 3G8 | 15 (48%) | 2 (6%) | 7 (23%) | 0 (0%) | 7 (23%) | 31 |
| 25 | | *enpp4* mRNA 2ng + cMO 20ng | 4A6 | 21 (68%) | 0 (0%) | 7 (23%) | 0 (0%) | 3 (9%) | 31 |
| 26 | S4C.D | *rfng* mRNA 2ng + *enpp4* MO2 10ng | 3G8 | 3 (20%) | 0 (0%) | 6 (40%) | 2(13%) | 4 (27%) | 15 |
| 27 | | *rfng* mRNA 2ng + *enpp4* MO2 10ng | 4A6 | 4 (27%) | 0 (0%) | 8 (53%) | 1 (7%) | 2 (13%) | 15 |
| 28 | S4E | *rfng* mRNA 2ng + cMO 10ng | 3G8 | 8 (38%) | 0 (0%) | 4 (19%) | 1 (5%) | 8 (38%) | 21 |
| 29 | | *rfng* mRNA 2ng + cMO 10ng | 4A6 | 9 (43%) | 0 (0%) | 8 (38%) | 0 (0%) | 4 (19%) | 21 |
| 30 | | *enpp4* MO2 10ng | 3G8 | 5 (28%) | 0 (0%) | 13 (72%) | 0 (0%) | 0 (0%) | 18 |
| 31 | | *enpp4* MO2 10ng | 4A6 | 6 (33%) | 0 (0%) | 12 (67%) | 0 (0%) | 0 (0%) | 18 |
| 32 | | *rfng* MO 20ng | 3G8 | 10 (53%) | 0 (0%) | 9 (48%) | 0 (0%) | 0 (0%) | 19 |
| 33 | | *rfng* MO 20ng | 4A6 | 16 (84%) | 0 (0%) | 3 (16%) | 0 (0%) | 0 (0%) | 19 |
| 34 | | *cMO 20ng* | 3G8 | 24 (78%) | 0 (0%) | 7 (22%) | 0 (0%) | 0 (0%) | 31 |
| 35 | | *cMO 20ng* | 4A6 | 24 (78%) | 0 (0%) | 7 (22%) | 0 (0%) | 0 (0%) | 31 |

**Supplementary Table 3B**

| Histological analysis compared | Figures compared | Injections compared | marker | Calculated p-value | Calculated p-value (with Bonferroni correction) | Standardized p-value (with Bonferroni correction) |
|---|---|---|---|---|---|---|
| 1/9 | 3A | *enpp4/cMO+LacZ* | *raldh1a2* | 1.04E-09 | 1.30E-07 | (0;0.001] |
| 2/11 | 3B | *enpp4/cMO+LacZ* | *rdh10* | 6.56E-08 | 8.20E-06 | (0;0.001] |
| 3/13 | 3C | *enpp4/cMO+LacZ* | *cyp26a1* | 2.42E-01 | 1.00E+00 | NS |
| 4/15 | 3D | *enpp4/cMO+LacZ* | *notch1* | 7.17E-15 | 8.96E-13 | (0;0.001] |
| 5/17 | 3E | *enpp4/cMO+LacZ* | *dll1* | 7.66E-12 | 9.57E-10 | (0;0.001] |
| 6/19 | 3F | *enpp4/cMO+LacZ* | *jag1* | 5.93E-14 | 7.41E-12 | (0;0.001] |
| 7/21 | 3G | *enpp4/cMO+LacZ* | *wnt4* | 8.93E-08 | 1.12E-05 | (0;0.001] |
| 8/9 | 3H | *enpp4MO/cMO* | *raldh1a2* | 1.21E-02 | 1.00E+00 | NS |
| 10/11 | 3I | *enpp4 MO/cMO* | *rdh10* | 1.34E-03 | 1.68E-01 | NS |
| 12/13 | 3J | *enpp4 MO/cMO* | *cyp26a1* | 4.71E-01 | 1.00E+00 | NS |
| 14/15 | 3K | *enpp4 MO/cMO* | *notch1* | 4.88E-02 | 1.00E+00 | NS |
| 16/17 | 3L | *enpp4 MO/cMO* | *dll1* | 4.00E-02 | 1.00E+00 | NS |
| 18/19 | 3M | *enpp4 MO/cMO* | *jag1* | 3.25E-04 | 4.06E-02 | (0.01;0.05] |
| 20/21 | 3N | *enpp4 MO/cMO* | *wnt4* | 1.29E-10 | 1.61E-08 | (0;0.001] |
| 22/24 | S4A/S4B | *rfngMO+enpp4RNA/ cMO+ enpp4 RNA* | 3G8 | 1.16E-01 | 1.00E+00 | NS |
| 23/25 | S4A/S4B | *rfngMO+enpp4RNA/ cMO+ enpp4 RNA* | 4A6 | 5.61E-01 | 1.00E+00 | NS |
| 26/28 | S4C.D/S4E | *enpp4MO2+rnfgRNA/ cMO+rfng RNA* | 3G8 | 3.52E-01 | 1.00E+00 | NS |
| 27/29 | S4C.D/S4E | *enpp4MO2+rnfgRNA/ cMO+rfng RNA* | 4A6 | 5.05E-01 | 1.00E+00 | NS |

**Supplementary Table 3. Scoring analysis of enpp4 mis-expression on RA. Notch and Wnt signalling pathways (related to Fig. 3 and Fig. S4). (S3A) Results from immunohistochemistry and *in situ* hybridization of *enpp4* mRNAs or MOs injected embryos.** Embryos were injected with *enpp4* RNA or MO alone or in combination with *rfng* RNA or MO. Embryos were scored for differences between the injected side (identified by Blue or Red Gal staining) and uninjected side acting as contra-lateral control side. The numbers of embryos displaying renal phenotype are indicated. The corresponding percentages are indicated in bracket. Each histological analysis was numbered as reference for the statistical comparisons in Supplementary Table 3B. Only the pronephros phenotype on the injected side is indicated. **(S3B) Statistical analyses.** Statistical comparisons between pairs of histological analysis listed in Supplementary Table 3A were carried out as indicated in the table. The Bonferroni multiple testing correction was applied to all Fisher's Exact Test. The calculated p value before and after correction and a standardized corrected p-value are given. NS: not significant

### Supplementary Table 4A

| Accession number | Full Protein name | Mw |
|---|---|---|
| Q9CQN1 | Heat shock protein 75 kDa. mitochondrial OS= *M.musculus* GN= Trap1 PE=1 SV=1 | 80158.65 |
| Q8BH59 | Calcium-binding mitochondrial carrier protein Aralar1 OS= *M.musculus* GN= Slc25a12 PE=1 SV=1 | 74522.94 |
| P38647 | Stress-70 protein. mitochondrial OS= *M.musculus* GN= Hspa9 PE=1 SV=2 | 73482.8 |
| Q62167 | ATP-dependent RNA helicase DDX3X OS= *M.musculus* GN= Ddx3x PE=1 SV=3 | 73056.16 |
| P63017 | Heat shock cognate 71 kDa protein OS= *M.musculus* GN= Hspa8 PE= 1 SV=1 | 70827.34 |
| P29341 | Polyadenylate-binding protein OS= *M.musculus* GN= Pabpc1 PE=1 SV=1 | 70598 |
| Q7TMK9 | Heterogeneous nuclear ribonucleoprotein OS= *M.musculus* GN= Syncrip PE=1 SV=2 | 69589.6 |
| Q8BGD9 | Eukaryotic translation initiation factor 4B OS= *M.musculus* GN= Eif4b PE=1 SV=1 | 68799.19 |
| Q91YQ5 | Dolichyl-diphosphooligosaccharide--protein glycosyltransferase subunit OS= *M.musculus* GN= Rpn1 PE=2 SV=1 | 68485.86 |
| P61222\| | ATP-binding cassette sub-family E member OS= *M.musculus* GN= Abce1 PE=2 SV=1 | 67271.14 |
| Q9Z0X1 | Apoptosis-inducing factor 1 OS= *M.musculus* GN= Aifm1 PE= 1 SV=1 | 66723.96 |
| O70194 | Eukaryotic translation initiation factor 3 subunit D OS= *M.musculus* GN= Eif3d PE= 1 SV=2 | 63948.5 |
| Q9Z247 | Peptidyl-prolyl cis-trans isomerase FKBP9 OS= *M.musculus* GN= Fkbp9 PE=1 SV=1 | 62955.57 |
| Q7TMK9-2 | Isoform 2 of Heterogeneous nuclear ribonucleoprotein Q OS= *M.musculus* GN= Syncrip | 62633.38 |
| P14685 | 26S proteasome non-ATPase regulatory subunit 3 OS= *M.musculus* GN= Psmd3 PE= 2 SV=2 | 60661.43 |
| P80317 | T-complex protein 1 subunit zeta OS= *M.musculus* GN= Cct6a PE=1 SV=3 | 57967.8 |
| Q6AX80 | Ectonucleotide pyrophosphatase/phosphodiesterase family member 4 OS= *X.laevis* GN= enpp4 PE= 2 SV=1 | 51278.25 |
| P61979 | Isoform 3 of Heterogeneous nuclear ribonucleoprotein K OS= *M.musculus* GN= Hnrnpk PE= 1 SV=1 | 50944.43 |
| Q99LC5 | Electron transfer flavoprotein subunit alpha. mitochondrial OS= *M.musculus* GN= Etfa PE=1 SV=2 | 34987.51 |

### Supplementary Table 4B

| Accession number | Full Protein name | Mw |
|---|---|---|
| Q9CQN1 | Heat shock protein 75 kDa. mitochondrial OS= *M.musculus* GN= Trap1 PE=1 SV=1 | 80158.65 |
| Q8BH59 | Calcium-binding mitochondrial carrier protein Aralar1 OS= *M.musculus* GN= Slc25a12 PE=1 SV=1 | 74522.94 |
| P38647 | Stress-70 protein. mitochondrial OS= *M.musculus* GN= Hspa9 PE=1 SV=2 | 73484.8 |
| Q62167 | ATP-dependent RNA helicase DDX3X OS= *M.musculus* GN= Ddx3x PE=1 SV=3 | 73056.16 |
| Q8K297 | Procollagen galactosyltransferase 1 OS= *M.musculus GN*= Glt25d1 PE=1 SV=3 | 71015.15 |
| P63017 | Heat shock cognate 71 kDa protein OS= *M.musculus GN*= Hspa8 PE=1 SV=2 | 70827.34 |
| Q99K51 | Plastin-3 OS= *M.musculus GN*= Pls3 PE=1 SV=3 | 70697.33 |
| P29341 | Polyadenylate-binding protein OS= *M.musculus* GN= Pabpc1 PE=1 SV=1 | 70598 |
| Q7TMK9 | Heterogeneous nuclear ribonucleoprotein OS= *M.musculus* GN= Syncrip PE=1 SV=2 | 69589.6 |
| Q91YQ5 | Dolichyl-diphosphooligosaccharide--protein glycosyltransferase subunit OS= *M.musculus GN*= Rpn1 PE=2 SV=1 | 68485.86 |
| P61222\| | ATP-binding cassette sub-family E member OS= *M.musculus* GN= Abce1 PE=2 SV=1 | 67271.14 |
| Q9Z0X1 | Apoptosis-inducing factor 1 OS= *M.musculus* GN= Aifm1 PE= 1 SV=1 | 66723.96 |
| O70194 | Eukaryotic translation initiation factor 3 subunit D OS= *M.musculus* GN= Eif3d PE= 1 SV=2 | 63948.5 |
| Q9Z247 | Peptidyl-prolyl cis-trans isomerase FKBP9 OS= *M.musculus* GN= Fkbp9 PE=1 SV=1 | 62955.57 |
| Q7TMK9-2 | Isoform 2 of Heterogeneous nuclear ribonucleoprotein Q OS= *M.musculus* GN= Syncrip | 62633.38 |
| P14685 | 26S proteasome non-ATPase regulatory subunit 3 OS= *M.musculus* GN= Psmd3 PE= 2 SV=2 | 60661.43 |
| P80317 | T-complex protein 1 subunit zeta OS= *M.musculus GN*= Cct6a PE=1 SV=3 | 57967.8 |
| P27773 | Protein disulfide-isomerase A3 OS= *M.musculus GN*= Pdia3 PE=1 SV=2 | 56642.78 |
| O70475 | UDP-glucose 6-dehydrogenase OS= *M.musculus GN*= Ugdh PE=1 SV=1 | 54797.35 |
| Q8R180 | ERO1-like protein alpha OS= *M.musculus GN*= Ero1l PE=1 SV=2 | 54050.26 |
| P97855 | Ras GTPase-activating protein-binding protein 1 member 4 OS= *M.musculus GN*= G3bp1 PE=1 SV=1 | 51796.96 |
| P61979 | Isoform 3 of Heterogeneous nuclear ribonucleoprotein K OS= *M.musculus* GN= Hnrnpk PE= 1 SV=1 | 50944.43 |
| Q99LC5 | Electron transfer flavoprotein subunit alpha. mitochondrial OS= *M.musculus* GN= Etfa PE=1 SV=2 | 34987.51 |

**Supplementary Table 4. Mass spectrometry analysis of membrane fractions (related to Fig. 4).** Proteins of control (A) or over-expressing *X.laevis* enpp4 protein (B) CHO cells membrane fraction were extracted and separated on a polyacrylamide gel. The band labelled by the Xlenpp4 antibody from the transfected Xlenpp4-pCDNA3 lane and the molecular weight equivalent band from the control lane were cut and analysed by mass spectrometry.

### Supplementary Table 5A

| | | | | Phenotypes | | | | | |
|---|---|---|---|---|---|---|---|---|---|
| Histological analysis | Figure | Injection | Marker stained | Normal | Enlarged | Reduced | Absent | Ectopic | Total number of scored embryos |
| 1 | 6A.B | *s1pr5* mRNA 2ng + *enpp4* mRNA 1ng | 3G8 | 29 (33%) | 2 (2%) | 23 (26%) | 1 (1%) | 34 (38%) | 89 |
| 2 | 6A.B | *s1pr5* mRNA 2ng + *enpp4* mRNA 1ng | 4A6 | 39 (44%) | 11 (12%) | 30 (34%) | 1 (1%) | 8 (9%) | 89 |
| 3 | 6C | *s1pr5* mRNA 2ng | 3G8 | 38 (73%) | 1 (2%) | 12 (23%) | 1 (2%) | 0 (0%) | 52 |
| 4 | 6C | *s1pr5* mRNA 2ng | 4A6 | 42 (81%) | 2 (4%) | 7 (13.5%) | 1 (2%) | 0 (0%) | 52 |
| 5 | 6D | *enpp4* mRNA 1ng | 3G8 | 64 (68%) | 9 (10%) | 14 (15%) | 0 (0%) | 7 (7%) | 94 |
| 6 | 6D | *enpp4* mRNA 1ng | 4A6 | 68 (72%) | 7 (7%) | 14 (15%) | 0 (0%) | 5 (5%) | 94 |
| 7 | S5A | *s1pr1* mRNA 2ng + *enpp4* mRNA 1ng | 3G8 | 57 (64%) | 12 (13.5%) | 13 (15%) | 0 (0%) | 7 (8%) | 89 |
| 8 | S5A | *s1pr1* mRNA 2ng + *enpp4* mRNA 1ng | 4A6 | 56 (63%) | 12 (13.5%) | 20 (22.5%) | 0 (0%) | 1 (1%) | 89 |
| 9 | S5B | *s1pr1* mRNA 2ng | 3G8 | 52 (90%) | 3 (5%) | 2 (3%) | 1 (2%) | 0 (0%) | 58 |
| 10 | S5B | *s1pr1* mRNA 2ng | 4A6 | 53 (91%) | 0 (0%) | 5 (9%) | 0 (0%) | 0 (0%) | 58 |
| 11 | S5C | *lpar1.1* mRNA 2ng + *enpp4* mRNA 1ng | 3G8 | 69 (65%) | 15 (14%) | 7 (7%) | 2 (2%) | 13(12%) | 106 |
| 12 | S5C | *lpar1.1* mRNA 2ng + *enpp4* mRNA 1ng | 4A6 | 81 (76%) | 12 (11%) | 7 (7%) | 2 (2%) | 4 (4%) | 106 |
| 13 | S5D | *lpar1.1* mRNA 2ng | 3G8 | 53 (96%) | 0 (0%) | 2 (4%) | 0 (0%) | 0 (0%) | 55 |
| 14 | S5D | *lpar1.1* mRNA 2ng | 4A6 | 53 (96%) | 0 (0%) | 2 (4%) | 0 (0%) | 0 (0%) | 55 |
| 15 | S5E | *p2yr10* mRNA 2ng + *enpp4* mRNA 1ng | 3G8 | 40 (78%) | 6 (12%) | 2 (4%) | 1 (2%) | 2 (4%) | 51 |
| 16 | S5E | *p2yr10* mRNA 2ng + *enpp4* mRNA 1ng | 4A6 | 45 (88%) | 3 (6%) | 1 (2%) | 2 (4%) | 0 (0%) | 51 |
| 17 | S5F | *p2yr10* mRNA 2ng | 3G8 | 49 (91%) | 2 (4%) | 3 (5%) | 0 (0%) | 0 (0%) | 54 |
| 18 | S5F | *p2yr10* mRNA 2ng | 4A6 | 53 (98%) | 0 (0%) | 1 (2%) | 0 (0%) | 0 (0%) | 54 |
| 19 | | *LacZ*mRNA 250pg | 3G8 | 54 (93%) | 0 (0%) | 4 (7%) | 0 (0%) | 0 (0%) | 58 |
| 20 | | *LacZ*mRNA 250pg | 4A6 | 55 (95%) | 0 (0%) | 3 (5%) | 0 (0%) | 0 (0%) | 58 |
| 21 | 6E.F | *s1pr5.L* MO 15ng | 3G8 | 24 (56%) | 2 (5%) | 16 (37%) | 0 (0%) | 1 (2%) | 43 |
| 22 | 6E.F | *s1pr5.L* MO 15ng | 4A6 | 17(39.5%) | 0 (0%) | 26(60.5%) | 0 (0%) | 0 (0%) | 43 |
| 23 | | cMO 15ng | 3G8 | 55 (90%) | 0 (0%) | 6 (10%) | 0 (0%) | 0 (0%) | 61 |
| 24 | | cMO 15ng | 4A6 | 53 (87%) | 0 (0%) | 8 (13%) | 0 (0%) | 0 (0%) | 61 |
| 25 | 6G.H | *s1pr5.L* MO 7.5ng + *enpp4* MO 5ng | 3G8 | 11 (26%) | 0 (0%) | 31 (74%) | 0 (0%) | 0 (0%) | 42 |
| 26 | 6G.H | *s1pr5.L* MO 7.5ng + *enpp4* MO 5ng | 4A6 | 8 (19%) | 0 (0%) | 34 (81%) | 0 (0%) | 0 (0%) | 42 |
| 27 | 6I | *s1pr5.L* MO 7.5ng + Control MO 5ng | 3G8 | 40 (78%) | 0 (0%) | 11 (22%) | 0 (0%) | 0 (0%) | 51 |
| 28 | 6I | *s1pr5.L* MO 7.5ng + Control MO 5ng | 4A6 | 36 (71%) | 0 (0%) | 15 (29%) | 0 (0%) | 0 (0%) | 51 |
| 29 | 6J | Control MO 7.5ng + *enpp4* MO 5ng | 3G8 | 14 (35%) | 0 (0%) | 26 (65%) | 0 (0%) | 0 (0%) | 40 |
| 30 | 6J | Control MO 7.5ng + *enpp4* MO 5ng | 4A6 | 16 (40%) | 0 (0%) | 24 (60%) | 0 (0%) | 0 (0%) | 40 |
| 31 | | cMO 12.5ng | 3G8 | 41(93.5%) | 2 (4.5%) | 1 (2%) | 0 (0%) | 0 (0%) | 44 |
| 32 | | cMO 12.5ng | 4A6 | 42 (96%) | 1 (2%) | 1 (2%) | 0 (0%) | 0 (0%) | 44 |
| 33 | 6K | *enpp4 2ng mRNA + s1pr5.L MO 15ng* | 3G8 | 18(47.5%) | 2 (7.5%) | 13(42.5%) | 0 (0%) | 7(17.5%) | 40 |
| 34 | | *enpp4 2ng mRNA + s1pr5.L MO 15ng* | 4A6 | 22 (55%) | 6 (15%) | 12 (30%) | 0 (0%) | 0 (0%) | 40 |
| 35 | | *s1pr5.L MO 15ng* | 3G8 | 25 (71%) | 2 (6%) | 6 (26%) | 0 (0%) | 0 (0%) | 35 |
| 36 | | *s1pr5.L MO 15ng* | 4A6 | 19 (57%) | 1 (3%) | 15 (45%) | 0 (0%) | 0 (0%) | 35 |
| 37 | 6L | *enpp4 2ng mRNA + cMO 15ng* | 3G8 | 13 (50%) | 6 (24%) | 7 (25%) | 0 (0%) | 24 (48%) | 50 |
| 38 | | *enpp4 2ng mRNA + cMO 15ng* | 4A6 | 26 (64%) | 7 (14%) | 12 (24%) | 0 (0%) | 5 (10%) | 50 |
| 39 | | *cMO 15ng* | 3G8 | 22 (76%) | 1 (3%) | 6 (21%) | 0 (0%) | 0 (0%) | 29 |

| Histological analysis | Figure | Injection | Marker stained | Phenotypes: Normal | Enlarged | Reduced | Absent | Ectopic | Total number of scored embryos |
|---|---|---|---|---|---|---|---|---|---|
| 40 | | *cMO 15ng* | 4A6 | 26 (90%) | 1 (3%) | 2 (7%) | 0 (0%) | 0 (0%) | 29 |
| 41 | S6E | *s1pr5.S MO 15ng* | 3G8 | 25 (38%) | 0 (0%) | 34 (51%) | 7 (11%) | 0 (0%) | 66 |
| 42 | | *s1pr5.S MO 15ng* | 4A6 | 26 (39%) | 0 (0%) | 19 (29%) | 21(32%) | 0 (0%) | 66 |
| 43 | | *s1pr5.L MO 15ng* | 3G8 | 12 (21%) | 5 (9%) | 28 (49%) | 12(21%) | 0 (0%) | 57 |
| 44 | | *s1pr5.L MO 15ng* | 4A6 | 22 (39%) | 0 (0%) | 9 (16%) | 26(45%) | 0 (0%) | 57 |
| 45 | | *cMO 15ng* | 3G8 | 48 (72%) | 2 (3%) | 14 (21%) | 3 (4%) | 0 (0%) | 67 |
| 46 | | *cMO 15ng* | 4A6 | 45 (67%) | 1 (1%) | 12 (18%) | 9 (14%) | 0 (0%) | 67 |

### Supplementary Table 5B

| Histological analysis compared | Figure compared | Injection compared | Marker | Calculated p-value | Calculated p-value (with Bonferroni correction) | Standardized p-value (with Bonferroni correction) |
|---|---|---|---|---|---|---|
| 1/5 | 6A.B/6D | *s1pr5 + enpp4/enpp4* | 3G8 | 1.08E-08 | 1.35E-06 | (0;0.001] |
| 2/6 | 6A.B/6D | *s1pr5 + enpp4/enpp4* | 4A6 | 1.50E-03 | 1.87E-01 | NS |
| 1/3 | 6A.B/6C | *s1pr5 + enpp4/s1pr5* | 3G8 | 8.80E-09 | 1.10E-06 | (0;0.001] |
| 2/4 | 6A.B/6C | *s1pr5.a + enpp4/s1pr5* | 4A6 | 1.15E-04 | 1.44E-02 | (0.01;0.05] |
| 3/19 | 6C | *s1pr5/ LacZ* | 3G8 | 7.66E-03 | 9.58E-01 | NS |
| 4/20 | 6C | *s1pr5/ LacZ* | 4A6 | 7.89E-02 | 1.00E+00 | NS |
| 7/5 | S5A/6D | *s1pr1 + enpp4/enpp4* | 3G8 | 8.64E-01 | 1.00E+00 | NS |
| 8/6 | S5A/6D | *s1pr1 + enpp4/enpp4* | 4A6 | 1.17E-01 | 1.00E+00 | NS |
| 7/9 | S5A/S5B | *s1pr1 + enpp4/s1pr1* | 3G8 | 1.32E-03 | 1.65E-01 | NS |
| 8/10 | S5A/S5B | *s1pr1 + enpp4/s1pr1* | 4A6 | 1.62E-04 | 2.03E-02 | (0.01;0.05] |
| 9/19 | S5B | *s1pr1/ LacZ* | 3G8 | 1.55E-01 | 1.00E+00 | NS |
| 10/20 | S5B | *s1pr1/LacZ* | 4A6 | 7.17E-01 | 1.00E+00 | NS |
| 11/5 | S5C/6D | *lpar1.1 + enpp4/enpp4* | 3G8 | 1.32E-01 | 1.00E+00 | NS |
| 12/6 | S5C/6D | *lpar1.1 + enpp4/enpp4* | 4A6 | 1.89E-01 | 1.00E+00 | NS |
| 11/13 | S5C/S6D | *lpar1.1 +enpp4/ lpar1.1* | 3G8 | 2.77E-05 | 3.47E-03 | (0.001;0.01] |
| 12/14 | S5C/S6D | *lpar1.1 + enpp4/ lpar1.1* | 4A6 | 8.39E-03 | 1.00E+00 | NS |
| 13/19 | S5D | *lpar1.1/LacZ* | 3G8 | 6.80E-01 | 1.00E+00 | NS |
| 14/20 | S5D | *lpar1.1/LacZ* | 4A6 | 1.00E+00 | 1.00E+00 | NS |
| 15/5 | S5E/6D | *p2y10 + enpp4/enpp4* | 3G8 | 1.28E-01 | 1.00E+00 | NS |
| 16/6 | S5E/6D | *p2y10 + enpp4/enpp4* | 4A6 | 6.73E-03 | 8.42E-01 | NS |
| 15/17 | S5E/S5F | *p2y10 + enpp4/p2y10* | 3G8 | 1.66E-01 | 1.00E+00 | NS |
| 16/18 | S5E/S5F | *p2y10 + enpp4/p2y10* | 4A6 | 8.43E-02 | 1.00E+00 | NS |
| 17/19 | S5F | *p2y10/ LacZ* | 3G8 | 4.05E-01 | 1.00E+00 | NS |
| 18/20 | S5F | *p2y10/ LacZ* | 4A6 | 6.19E-01 | 1.00E+00 | NS |
| 21/23 | 6E.F | *s1pr5.L*MO/cMO | 3G8 | 1.22E-04 | 1.52E-02 | (0.01;0.05] |
| 22/24 | 6E.F | *s1pr5.L*MO/ cMO | 4A6 | 5.39E-07 | 6.74E-05 | (0;0.001] |
| 41/45 | S6E | *s1pr5.S*MO/ cMO | 3G8 | 8.61E-05 | 1.08E-02 | (0.01;0.05] |
| 42/46 | S6E | *s1pr5.S*MO/ cMO | 4A6 | 3.22E-03 | 4.02E-01 | NS |
| 41/43 | S6E/6F | *s1pr5.S*MO/ *s1pr5.L*MO | 3G8 | 1.08E-02 | 1.00E+00 | NS |
| 42/44 | S6E/6F | *s1pr5.S*MO/ *s1pr5.L*MO | 4A6 | 1.57E-01 | 1.00E+00 | NS |
| 25/27 | 6G.H/6I | *s1pr5.L*MO + *enpp4*MO/ *s1pr5.L* MO + cMO | 3G8 | 8.21E-07 | 1.03E-04 | (0;0.001] |
| 26/28 | 6G.H/6I | *s1pr5.L*MO + *enpp4*MO/ / *s1pr5L* MO + cMO | 4A6 | 9.37E-07 | 1.17E-04 | (0;0.001] |
| 25/29 | 6G.H/6J | *s1pr5.L*MO + *enpp4*MO/*enpp4* MO + cMO | 3G8 | 4.74E-01 | 1.00E+00 | NS |
| 26/30 | 6G.H/6J | *s1pr5.L*MO + *enpp4*MO/*enpp4* MO + cMO | 4A6 | 5.21E-02 | 1.00E+00 | NS |
| 33/37 | 6K/6L | *enpp4RNA + s1pr5.LMO /enpp4RNA* + cMO | 3G8 | 4.18E-03 | 5.23E-01 | NS |
| 34/38 | 6K/6L | *enpp4RNA + s1pr5.LMO /enpp4RNA* + cMO | 4A6 | 2.37E-01 | 1.00E+00 | NS |

**Supplementary Table 5. Scoring analyses of enpp4 and lipidic receptors mis-expression (related to Fig. 6. And Fig.S6). (A) Results from immunohistochemistry of enpp4 and lipidic receptors mRNA and MO injected embryos.** Embryos were either injected with *s1pr1*. *s1pr5*. *lpar1.1*. *p2y10* and *enpp4* mRNA alone or in combination or with *s1pr5.L s1pr5.s* and *enpp4* MO alone or in combination. Embryos were scored for differences between the injected side (identified by Blue Gal staining) and uninjected side acting as contra-lateral control side. The numbers of embryos displaying renal phenotype are indicated. The corresponding percentages are indicated in bracket. Each histological analysis was numbered as reference for the statistical comparisons in Supplementary Table 5B. Only the pronephros phenotype on the injected side is indicated. **(B) Statistical analyses.** Statistical comparisons between pairs of histological analysis listed in Supplementary Table 5A were carried out as indicated in the table. The Bonferroni multiple testing correction was applied to all Fisher's Exact Test. The calculated p value before and after correction and a standardized corrected p-value are given. NS: not significant.
